# The Impact of Grant Funding on the Publication Activity of Awarded Applicants: A Systematic Review of Comparative Studies and Meta-analytical Estimates

**DOI:** 10.1101/354662

**Authors:** Ruslan T. Saygitov

## Abstract

The connection between grant funding and research productivity has not been well established.

**Objective:** to examine the impact of grant funding on the publication activity of awarded applicants.

**Methods:** a systematic review of results from comparative studies on the publication activity of applicants (awarded vs rejected) both prior to and after the award process. All pool estimates (weighted mean difference) were based on random-effects models.

**Results:** revealed 16 relevant publications (grant funding from 14 funds, 1980 to 2007 years), all with results from quasi-experimental studies. 45 paired values (ex ante – ex post) for the number of articles published by awarded and rejected applicants were used in the quantitative synthesis. The median average publication activity of awarded applicants before the award process was 2.4 (1.3; 3.4) and after the award process 3.1 (1.7; 4.3) publications per year, for rejected applicants was 1.8 (1.0; 2.9) and 2.4 (1.1; 3.8) respectively. The summation of the results from these studies using the difference-in-differences approach showed that awarded applicants published 0.14 articles per year (95% CI 0.07 to 0.21) more than rejected applicants (adjusted for publication bias). A meta-regression analysis made it possible to tie together the revealed small difference with the difference-in-differences approach bias − the subsequent differences in the groups are determined by the scale of the initial differences in their publication activity.

**Conclusion:** awarded applicants published slightly more often than their rejected opposites. However, this effect may be the result of a bias caused by the shortcomings of the difference-in-differences approach.

## BACKGROUND

Funding for global research in recent years has remained at 1.8% of gross domestic product, or just over 1.5 trillion dollars in nominal terms (of this roughly 80% of all research spending is by the 10 global leaders in R&D) [1], A significant proportion of these funds (roughly 20-40%, according to some estimates) is distributed in the form of grants [2], The positive impact of scientific funding on research productivity would seem to be obvious and is corroborated by macroeconomic assessments [3, 4], However, the effectiveness of individual funding mechanisms (basic, competitive, contract, subsidized) and support (tax) for research activities remains surprisingly hypothetical.

The effectiveness of grant funding has been studied for several decades. The very first study on this subject showed that the number of articles published after a competitive award process by funded applicants far surpassed the number published by those who were not funded [5]. However, the design of research studying the impact of grants on the research productivity of funded applicants is extremely diverse. They include comparative and non-comparative (only funded applicants) studies, ‘before and after’ or only ‘after’ studies, and studies on ‘dose’-dependent effects (where ‘dose’ is amount of money). In comparative studies the control group might be non-funded applicants or colleagues not participating in the competitive award process. Furthermore, they may also compare the research productivity not only of individual researchers, but also groups of researchers (laboratories, research centres, etc.). The diversity in research design is further amplified by the variety in the tested effects and corresponding indicators of research productivity. According to several systematic estimates, the number of such indicators is over ten [6, 7], However, research productivity is most frequently appraised using bibliometric indicators, in particular the Hirsch index, the number of works published and citations [7].

A systematic summary of the results of studies on the effectiveness of grant funding has been carried out before in the context of (bio)medical research [8]. The review compared the publication activity of awarded and rejected applicants in quasi-experimental studies using a ‘before and after’ design. It showed that funded applicants published on average one paper more than their opposites in the 4-5 years after the competitive award process. It did however reveal that a ‘treatment’ effect was caused by the different initial publication activity of the applicants (awarded vs. rejected) [8]. Whether the displacement detected in estimates of grants’ impact on the publication activity of awarded applicants is typical of the (bio)medical field or universal requires further study.

### Research objective

To examine the impact of grants on the publication activity of awarded applicants based on a systematic analysis of data from comparative studies (awarded vs. rejected) and the results of meta-analytical estimates which take into account the initial differences in the applicants.

## METHODS

The research planning and description of the search procedures and results were all carried out in line with recommendations [9].

### Eligibility criteria

*Types of studies:* controlled studies (prospective or retrospective, randomized or quasi-experimental).

*Types of participants (controls):* awarded and rejected applicants (not populations or colleagues for control group; not awarded applicants in other competitive award processes; not unit or research group analysis).

*Types of intervention (‘treatment’):* research, training or other types of grants (non-repayable funds for individuals); grant award conditions – completions through peer-review.

*Types of outcome measures:* number of articles published (*ex-ante* or before and *ex-post* or after competitions).

No publication date, publication status (working paper, draft version of article, unpublished material, and abstracts) or funding duration restrictions were imposed.

### Information sources

Studies were sought using Google Scholar (http://scholar.google.ru/) and Google Search (http://google.ru). Both search systems were used in parallel, either duplicating one another (when searching for key words) or individually (Google Scholar when search for articles from reference lists, citations and related publications; Google Search when manually searching for data only published via Internet resources).

### Search strategy

Before starting the systematic search, the author had a collection of articles (>100) on the topic in question. The collection of articles was created by means of a manual cross (but not systematic) search using Google Search conducted between November 2012 and May 2013. The objective of the search was to analyse the literature to justify the research, the results of which have already been published [8]. Later, to compare the Russian data with the estimate of grants’ impact on research productivity in other countries, in August-December 2013 a systematic literature analysis was carried out.

The systematic search was carried out using the ‘snowball’ technique. The search comprised three stages. The **first stage** involved searching for key words using search filters (in Google Scholar and Google Search). Three main key words had to be present in the text of the publications: ‘grant’, ‘productivity’ and ‘bibliometric’. In addition to this, at least one of the following key words had to be present in the text of the publications: ‘unfunded’ OR ‘unsuccessful’ OR ‘rejected’. The assumption was that the main key words would ensure a higher degree of sensitivity in the research (i.e. access to a larger number of relevant sources), while the additional key words would allow greater specificity (i.e. they would exclude from the search sources containing research results without the appropriate control groups). The first 500 links were examined from Google Scholar and the same from Google Search.

The links were copied into an MS Excel table, sorted by the name of the cited document, and duplicate links were then removed from the analysis. The names of the remaining publications were studied, together with brief summaries of the content where applicable. Using this information, a list of articles was compiled for further study of the complete text.

The **second stage** of the search (using Google Scholar only) involved reviewing i) brief summaries of the material (articles, etc.) content from the reference lists of relevant publications (from the first stage of the search), ii) publications citing relevant publications, and iii) the first 50 related publications (links to citations and related articles are automatically created by Google Scholar; links in Arabic and Chinese were excluded). When relevant publications were found (those found in the first stage were not considered), the publications from the reference list citing related articles were examined.

The **third stage** of the search comprised a content analysis of the publications identified ‘by chance’ in the previous stages, i.e. not through the lists of cited, citing or related articles. Information on the name and/or brief description of the content of publications arousing interest was kept in a separate file and analysed after the key words and related sources search (references, citations and related articles list). When relevant publications were identified (those identified in the previous stages were not taken into account), the search for related sources was repeated.

### Data collection process

From the articles meeting the search criteria, digital data was extracted on the total number of published articles or the average number of articles per year. Similar steps were taken with regard to the number of citations. For the systematic review, assessments of applicants’ publication activity in certain fields or summary data in multi-disciplinary competitive award processes were used.

The translation of key fragments (for example, the captions of rows and columns in tables) in non-English articles (original text languages included German and Danish) into English was carried out using the online Google Translate (https://translate.google.ru).

The numerical data that had only been presented graphically in the published paper was digitized using GetData Graph Digitizer (http://getdata-graph-digitizer.com/).

When duplicate publications were detected, the source with the more complete information was taken into account (and then cited). If the data required for the systematic review and meta-analytical estimate was identical in duplicate publications (in particular, identical samples and outcomes), the free access publication or the article published in a scientific journal was cited. When discrepancies in the published data were detected (different samples and/or outcomes), the data published in peer-reviewed journals were cited (if duplicates were only published in journals, data from the journal with the highest impact factor were cited).

A. Schubert presented individual data on the publication activity of applicants [10]. The analysis of these data showed that the bibliometric data for awarded applicants included publications sampled over a longer period (on average +1 year). As a result, groups of awarded and rejected applicants (only principal researchers^1^ matched to take into account the year of the competitive award process were formulated from the individual data.

In B. Lal et al., the data presented did not allow for an evaluation of publication activity over equal time frames before and after the competitive award process [11]. Using the list of applicants (awarded and rejected) provided by the authors, the number of published works was re-calculated (with a time window of 7 years before and after the process) using data from the Scopus database (http://www.scopus.com). Works published in peer-reviewed journals were taken into account.

### Data items

Information was extracted from each included study (where available) on: (1) the characteristics of applicants (including age, education); (2) the characteristics of the competitions (country and year, fields, grant type, average funding amount); (3) the characteristics of the study (including the design, inclusion and exclusion criteria, matching and methods); (4) bibliometric data (including the source of the data, the period of the data before and after the competitive process, the number of published articles and citations, the impact factor of the journals in which the articles are published); and (5) other outcomes (subsequent grant funding, career growth).

### Summary measures

The primary outcome measure was the number of published articles (per year per person). The effect size relating to the grant funding effect for a single study was determined using the difference-in-differences approach (the difference between the pre-post number of articles in the ‘treatment’ and control groups). The difference in the groups was adjusted using the initial publication activity of applicants (before the competitive award process).

### Planned methods of analysis

The meta-analyses were performed by computing the pooled effect size using random-effects model. The pooled estimate (weighted mean difference, WMD) and 95% confidence intervals (CI) were calculated. Between-study variability was tested by measure of inconsistency (I^2^, the percentage of variability in effect size across studies due to heterogeneity rather than chance). Publication bias was evaluated using a funnel plot of the study effect size by the inverse of log(sample size). The asymmetry of the funnel plot was formally evaluated with Egger’s test. Meta-analytical estimates were carried out using the StatsDirect statistical package (http://www.statsdirect.com). The meta-regression analysis was carried out using linear regression (IBM SPSS Statistics v. 22) and the beta value (standard error, SE) was calculated, together with the explained dispersion of the dependent variable (adjusted R squared).

## RESULTS

### Study selection

A detailed description of the results of the systematic search is given in **Appendix A.** During the search the titles and abstracts of 1,979 non-duplicate publications (65% of 3,034 publications found in the search) and then 192 (10%) full texts were briefly analysed. A total of 16 publications were identified for inclusion in the review [8, 10,11, 12-24].

### Study characteristics

A detailed description of the 16 relevant studies is given in **Appendix B.** All of the studies included in the systematic review are quasi-experimental retrospective studies with regard to the selection of participants and prospective outcome (publication activity) measures. No randomized studies on the effectiveness of grant funding were found. 13 studies were published in English, 2 in Danish [19, 20], and 1 in German [16]. The results from 7 of the 16 studies were published in peer-reviewed journals (the remainder–grant funding agencies reports and working papers – were made available online).

From the studies found, the results of funding from 14 funds in 10 countries were analysed, and one study presented combined data on EU applicants [18]. The studies covered funding programmes during the period 1980 to 2007.

The bibliometric analysis in all of the studies included publications in scientific (peer-reviewed) journals. In addition, 2 studies [17, 20] took into account conference papers (proceedings) and one [20] – books. Data in 4 studies [16, 18, 20, 23] were broken down by scientific field and another 4 [8, 17, 18 (EMBO data set), 24] by groups based on applicants scientific qualifications or participants in different competitive award processes. In total, 45 paired values (ex ante – ex post) for the number of articles published by awarded and rejected applicants were used in the quantitative synthesis. The median number (25th; 75th quartiles) of applicants in groups (n=45) was 40 (20; 112) for awarded applicants and 71 (38; 192) for rejected applicants.

The duration of the periods covered by the bibliometric data before and after the competitive award process ranged from 3 to 7 years. The majority of the groups (22 of 45; 49%) took into account publication activity over 5 years before and after the award process [8, 15, 19-23]. 4-year (16/45; 36%) [12-14, 16, 17, 24] and 3-year periods (6/45; 13%) [10, 18] were analysed less frequently. One study analysed the publication activity of applicants in the 7 years before and after the award process [11].

In some studies the average publication activity of awarded applicants before the award process varied from 0.1 to 7.1 publications per year, with a median of 2.4 (1.3; 3.4), and after the process from 0.1 to 7.6 publications per year, with a median of 3.1 (1.7; 4.3). The average publication activity of rejected applicants before the award process varied from 0.1 to 5.6 publications per year, with a median of 1.8 (1.0; 2.9), and after the process from 0.1 to 5.7 publications per year, with a median of 2.4 (1.1; 3.8) (details see in **Appendix C**).

### Syntheses of results

The pooled WMD for the change in publication activity of awarded and rejected applicants after the competitive award process based on the data of the 16 studies (45 groups, 48 825 awarded and 28 203 rejected applicants) was 0.17 publications per year (95% CI 0.10 to 0.24). After adjustment for publication bias^2^ (for more detail see **Appendix D**) by excluding from the analysis the data of 4 groups with marginal data [10, 11, 16 (Physics data set), 20 (Medicine data set)] (all published in non-peer reviewed sources), the pooled WMD value was 0.14 (95% CI 0.07 to 0.21). However, generally, and after adjustment to take into account publication bias, the results of the studies have been found to exhibit significant statistical heterogeneity (I^2^ >60%). The heterogeneity was not able to be adjusted taking into account the mean age of the award process participants (<= 40 years/other [including groups with unknown ages of applicants]), fields ([bio]medicine/other), and source of data (peer reviewed journal/other).

### Additional analyses

The analysis of the paired values shows that the effect size was dependent on the scale of the differences in the comparable groups (awarded vs rejected) before the competitive award process: the higher the initial difference in average publication activity values the higher the mean effect size recorded subsequently (Fig. 1). In addition, it was also noted that differences in the comparable groups could be affected by awarded applicants with the highest publication activity (Fig. 2). If we exclude from the analysis applications with ≥ 4 publications per year (awarded and/or rejected), the differences between the comparable groups cannot be seen: for the whole dataset (45 paired groups), the baseline publication adjusted beta = 0.24 (SE 0.11) (p=0.035), and when excluded (38 paired groups), the beta = 0. 14 (SE 0.11) (p = 0.214).

Excluding the results of the biggest studies [21, 22] from the analysis did not have any impact on the scale of the differences – the pooled WMD was 0.18 (95% CI 0.09 to 0.28). The estimated effect in studies published in scientific journals is near zero (pooled WMD = 0.05 (95% CI -0.07 to 0.08)), while the results of studies published in other (non-peer-reviewed) sources showed a higher publication activity among awarded applicants (pooled WMD = 0.26 (95% CI 0.15 to 0.37)).

**Figure 1.**
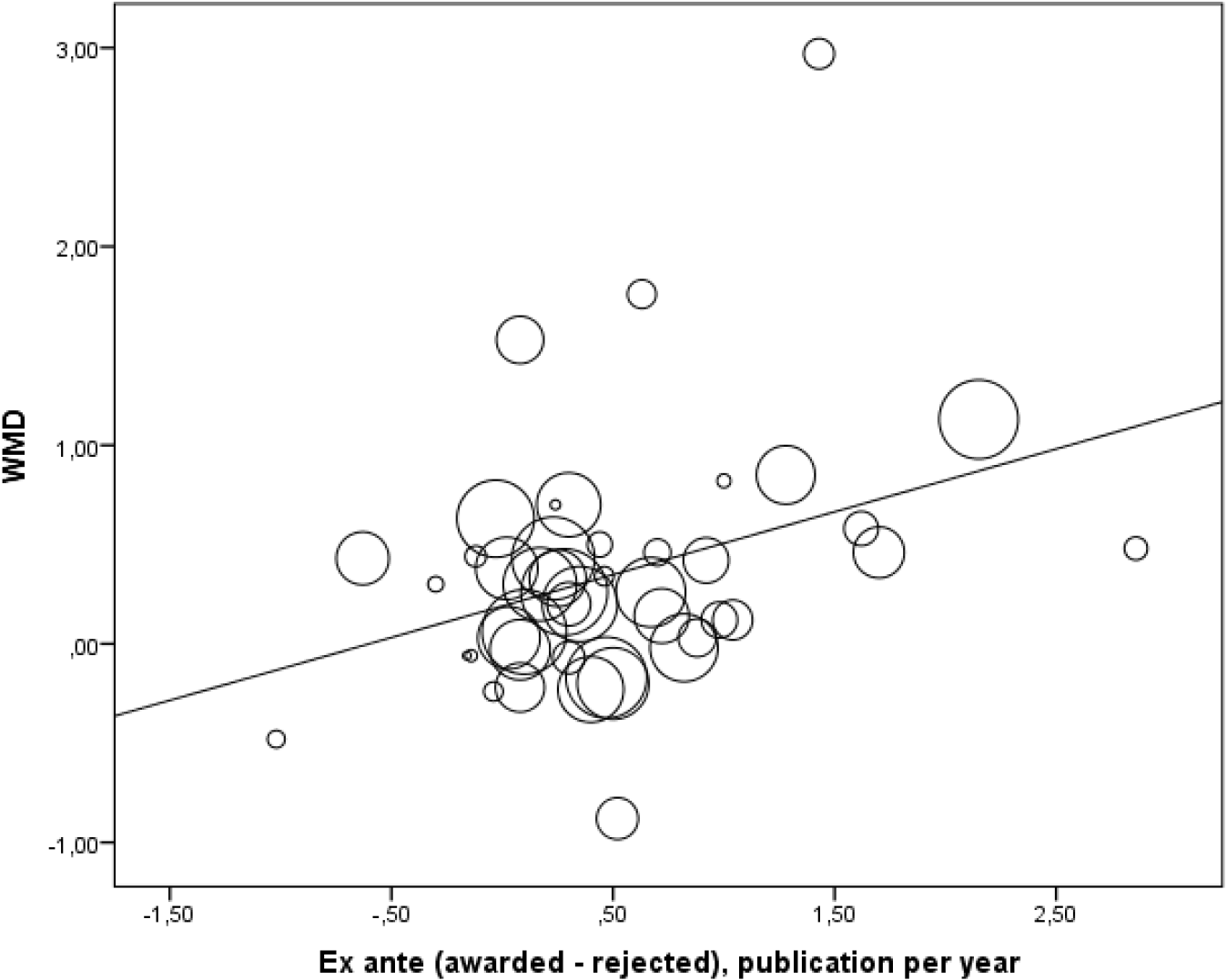
The effect size of grant funding is dependent on the initial differences between the groups of awarded and rejected applicants

**Note.** The size of the circle is a logarithm of the total number of applicants (awarded and rejected), beta = 0.32 (SE 0.13) (p = 0.016); R2 = 0.22. Solid line = regression line.

**Figure 2.**
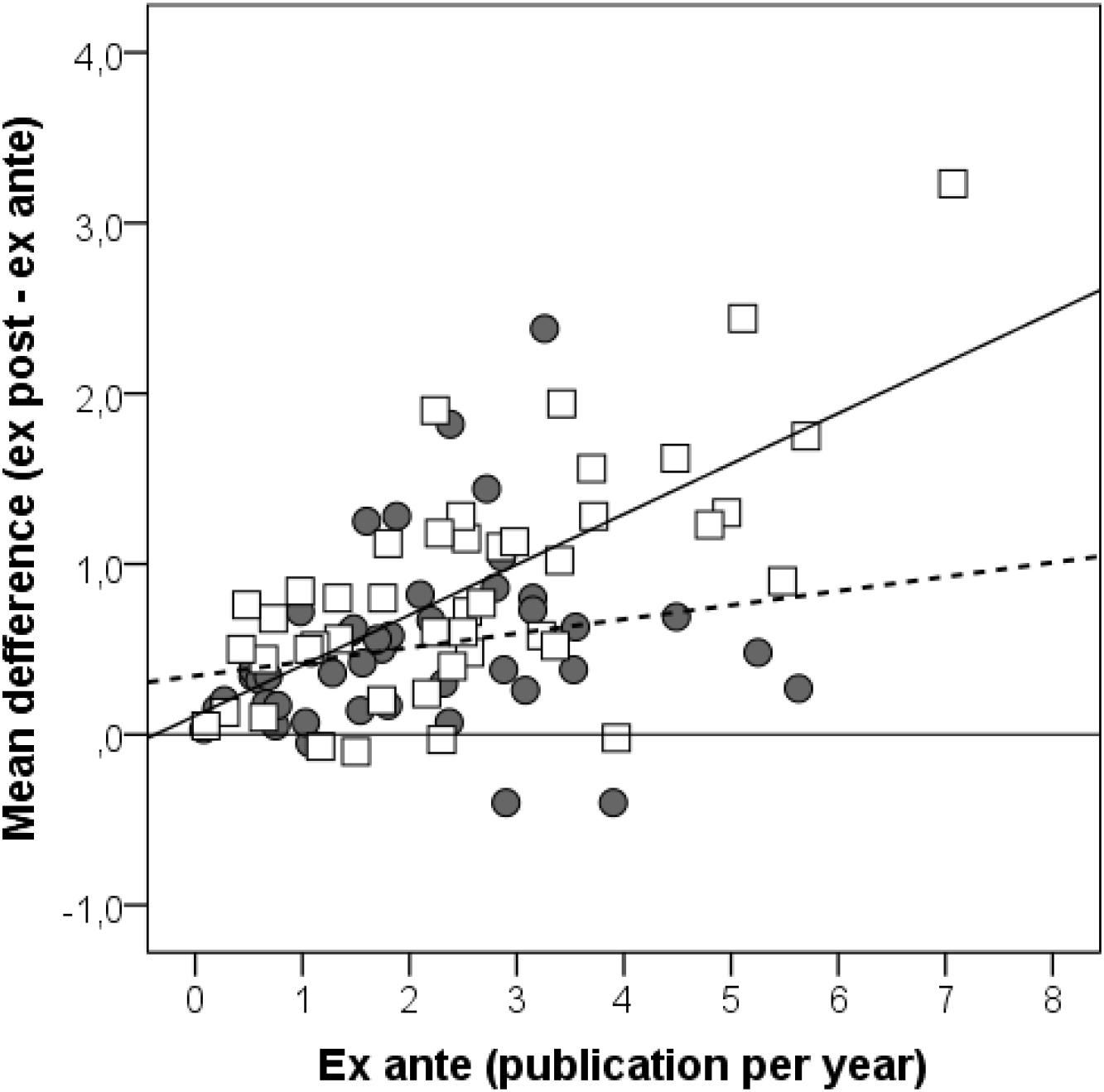
The change in publication activity after the award process in awarded and rejected applicants with different initial publication activity: results from a meta-regression analysis

**Note.** Square markers are awarded applicants; circular markers are rejected applicants. Solid line = regression line for the awarded group; dashed line = regression line for rejected applicants.

## DISCUSSION

### Summary of the main results

The systematic review and meta-analytical estimates of the results of quasi-experimental studies have showed a small (roughly +1 publication over 6 years, 95% CI 4 to 10 years) increase in publication activity linked to grant funding for awarded applicants. However, a publication bias was detected, caused by the results of studies published in non-peer-reviewed sources (unpublished data, grant funding agencies reports, working papers). After adjusting the meta-estimates of publication activity by excluding the data causing the bias from the analysis, the relationship between grant funding and the increase in publication activity became even more modest (roughly +1 publication over 7 years, 95% CI 5 to 14 years).

### ‘Treatment’ effect or approach bias?

There are grounds for arguing that the small ‘treatment’ effect is the result of a bias [25]. In turn, we can suggest that even the small increase in publication activity among awarded applicants, revealed in the meta-analysis, was caused not by grants, but the estimation method. A DID approach is often used to detect the effect of interference in non-randomized comparative studies. This approach takes into account the baseline values of dependent variables, but not the size of the differences in these values. The meta-regression analysis (see Fig. 1) showed that the scale of these differences is a significant variable, i.e. the higher the initial difference in publication activity between awarded and rejected applicants, the higher the effect of the grants (pooled WMD) will have^3^. And vice versa, the change in publication activity among awarded and rejected applicants with the same initial values was the same (see Fig. 2). The results of matched-pair case-control studies (awarded and rejected applicants with the same initial publication activity) confirm this assessment [14, 18, 20].

The question about impact of grant funding on publication activity of awarded applicants with marginal initial publication activity (≥ 4 publications per year) remains open. It should be noted that applicants with the highest (i.e. rare) publication activity are generally excluded from making awarded-rejected pairs in matched case-control studies. When the whole cohort is analysed, such applicants would cause a DID-bias described above. Furthermore, due to lack of comparable initial publication activity values for the control group, regression analysis would also result in overrating the impact of grant funding. For this reason, the results of the regression analysis may differ from the results of matched-paired case-control studies, and show a small positive effect of grant funding on applicants’ publication activity [18]. A similar result was also obtained after conducting this systemic review (see **Additional analyses**).

Thus the available data indicates that DID-based approach and regression analysis tend to overestimate grants’ impact on publication activity of awarded applicants. This bias is levelled by adjusting the results taking into account initial differences between awarded and rejected applicants’ key predictor values affecting their subsequent publication activity – initial (past) number of published papers [26-28].

### Publication activity as a funding effect indicator

Sustainable development of human potential, society, and environment are among the ultimate goals of R&D funding and grant funding in particular (‘hard’ outcome). Therefore progress in accomplishing these goals should be taken into account when assessing funding effects. However, this would only be possible on macro-level (for specific countries, or the whole world). On micro-level (e.g. individual researchers or research units), funding effects can be evaluated by measuring research productivity: output and impact [29]. There’s no doubt that researchers’ publication activity, along with their patent activity, conference presentations and shared databases, constitutes a research output [29].

Furthermore, we know that researchers’ publication activity also serves as a predictor of scientific value of their future work [30-32], Thus there are grounds to believe that the number of published papers is a proxy of research productivity, individual and aggregated (on micro-level). Furthermore, certain studies demonstrate that the number of published academic papers is a proxy of research impact on ‘hard’ outcome on macro-level too [33].

The systemic analysis did not reveal any impact of grants on publication activity of awarded applicants (vs rejected). Does that mean that research grants (and this was the kind of grants whose effect was analysed in 14 out of 16 studies included in the systemic review) were a waste of public money? To confidently answer this question we must study the impact of grant funding on other aspects of research productivity. Strictly speaking, the results of the systemic review do not extend to other characteristics of research productivity, which are measured using other nominal or normalised output/impact metrics. However, the analysis of individual data on citations obtained for NDPA applicants (**Appendix E**) [11] suggests that the effect of grants in this case is due to the initial differences between awarded and rejected applicants. Similar conclusions can also be reached by analysing normalised publication activity [13] and normalised citations [34], There is no conclusive evidence of research grants’ impact on awarded applicants’ patent activity either (vs rejected) (see, e.g., [13]). Note that a grant application always includes a specific research plan clearly indicating the study’s objectives, steps to be taken to accomplish them, and expected results. So if the study is completed its results must be published, even if they happen to be negative or have been patented beforehand. Hiding such data obviously constitutes an example of less-than-optimal application of public resources allocated to fund R&D.

Of course one could argue that publications (just as other kinds of research outputs) are not the goal of grant funding agencies’ operations. However, I believe we shouldn’t forget about researchers. Did high publication activity cease to bring rewards, obvious and hidden ones? Did the principle ‘publish or perish’ lose relevance? Obviously publications were, and remain, one of the most important research outputs, and a major motivation factor for researchers. Furthermore, for young researchers high publication activity can turn into an “evolutionary” advantage [35]. Whether it’s good or bad, is a topic for another discussion.

Finally, there are grounds to believe that estimates of grant funding impact on research productivity can be underrated due to the presence of a competitive subsidy market for scientific research, where rejected applicants can obtain comparable (vs awarded ones) funding after the award process (discussed in [22]). However, studies show that the opposite is true: awarded applicants tend to attract additional funding from other sources [13].

### Adequate proof of funding effects

Firm evidence of the impact of competitive grants funding on research productivity (output and impact) of applicants should be found in randomised studies. This requires testing hypotheses regarding grants’ *efficacy* (effect of money only), *effectiveness* (effect of money allocated through a competitive award process which ensures that “the best” are rewarded), and *efficiency* (effect of money allocated through a competitive award process taking into account tender-related expenditures and waste of money due to inadequate choice of “the best”).

The first randomised study of grants’ impact on research productivity is currently under way [36]. It analyses the effect of money only (the grant efficacy): applicants will be randomly assigned into ‘treatment’ or control groups (the results are expected to be obtained in 2020 at the earliest). However, the study will not answer the question about the effect of money in the presence of the competitive award process including application of the existing peer-review system; inefficiency of this system is subject of extensive discussions (examples include [18], however, the literature is more extensive).

The hypothesis testing of the effect of money in the conditions of a real award process call for a randomised 2×2 factorial experiment, in which the experimental groups (‘treatment’ or control) will be formed within field-specific awarded and rejected groups (with stratification for the size of the requested funding and previous research productivity). One would expect that in an effective competitive award process the effect of money will be apparent in the group of awarded applicants (‘treatment’ vs control) or, alternatively, it will be significantly higher in the awarded group than in the rejected group (all – those who have received the funding sought).

## LIMITATIONS

Meta-analysis results are considered in the framework of two hypotheses: the results cover the whole population (common effect size) or constitute the mean of all true effects [37], This systemic review is based on the hypothesis with the mean of all true effects at the core. However, both these hypotheses need further testing. Which means it remains unclear whether the obtained meta-analytic estimates hold true for grant-based research funding in general, or only for specific fields or subfields.

There’s a probability that meta-analytic estimates produced were incorrect due to biases, known and hypothetical ones [37, 38]. Some of the more common biases (publication and approach bias) were studied. These data increases confidence in the presented results, but does not completely exclude possibility of error, e.g. due to the studies’ incompatibility in terms of grant size. Keeping in mind that most of the studies included in the systemic review lack such data, the meta-analytic estimates have not been adjusted on the basis of grant size. We believe that the very fact of grant allocation provided sufficient grounds to compare awarded and rejected applicants’ performance. Which, however, does not exclude the need to assess ‘dose’-dependent effects.

The study results were found to be statistically heterogeneous (I^2^ ≥60%), which cannot be explained taking into account the available independent variables (applicants’ age, fields, data sources), or when adjusting for publication bias. A random-effects model is recommended for analysing data in such situations [9]. Furthermore, we suggest to assume that “statistical heterogeneity is inevitable. … Heterogeneity will always exist whether or not we happen to be able to detect it using a statistical test.’.[9].

The search for research studies for this systematic review was mainly conducted by way of Google-indexed webpages. It could, however, be argued that searching across just single database may be imperfect [39]. That being said, it has been demonstrated quite convincingly in modeling a real systematic search that the Google search engine is a tool that retrieves over 90% of all relevant publications from Google’s “database” [40, 41]. It is important to note that data collection and search were carried out by one researcher. For some data, participation of at least two researchers may reduce the possibility of rejecting relevant reports by 8% [42],

## CONCLUSION

Is there a benefit from grant funding? This systemic review has produced evidence suggesting a negative answer to this question regarding publication activity of awarded applicants. However, common sense implies grant money should produce at least some effect. Why no such effects can be confirmed? Possibly the reason is simple: frequently grant funding does not involve an obligation to produce publications, patents, or other R&D outputs. Does this lead to missing research results (albeit negative ones), and thus inefficient application of resources the society allocates to fund R&D? On the other hand, funding effects may be levelled by problems with evaluation of grant applications. If we accept this assumption, we would have to abandon the complex and costly peer-review system in favour of allocating grants based on panels of objective research activity indicators and available infrastructure, prior to competitive award process. Answering these difficult questions requires additional studies. Their results would extend our understanding of grant funding efficiency, whose benefits should be analysed under more strict experimental conditions.

## Acknowledgments

I am indebted to Callum Walker and Vsevolod Korolev for their help in translating this article. I would also like to thank Galina Kitova and Stanislav Zaichenko (both from Higher School of Economics, Moscow) for their constructive criticism discussion of the manuscript.

## Footnotes

1 The individual bibliometric data on supported projects contained information on principal researchers and other participants (for unsupported projects, the results were only based on the publication activity of the principal researchers).

2 Egger test: bias = 0.75 (95% CI -0.17 to 1.66), p = 0.106

3 Consider 3 scenarios: the initial publication activity of rejected applicants is 1, 2 or 3 publications per year and awarded applicants 2, 4 or 6 respectively. Thus the initial difference in publication activity equal to 1, 2 and 3. If initial differences do not matter, expected impact of the competitive award process for all of these scenarios will be the same (suppose DID values will be 1). However, we observed (see Fig. 1) that DID values are 1, 1.32 and 1.64 for the different scenarios respectively. Given equal initial publication activity of rejected and awarded applicants, the above bias is levelled (see Fig. 2).

